# Evaluating test-retest reliability and sex/age-related effects on temporal clustering coefficient of dynamic functional brain networks

**DOI:** 10.1101/2021.10.21.465376

**Authors:** Yicheng Long, Chaogan Yan, Zhipeng Wu, Xiaojun Huang, Hengyi Cao, Zhening Liu, Lena Palaniyappan

## Abstract

The multilayer dynamic network model has been proposed as an effective method to understand how the brain functions dynamically. Specially, derived from the definition of clustering coefficient in static networks, the temporal clustering coefficient provides a direct measure of topological stability of dynamic brain networks and shows potential in predicting altered brain functions in both normal and pathological conditions. However, test–retest reliability and demographic-related effects on this measure remain to be evaluated. Using a publicly available dataset from the Human Connectome Project consisting of 337 young healthy adults (157 males/180 females; 22 to 37 years old), the present study investigated: (1) the test-retest reliability of temporal clustering coefficient across four repeated resting-state functional magnetic resonance imaging scans as measured by intraclass correlation coefficient (ICC); and (2) sex- and age-related effects on temporal clustering coefficient. The results showed that (1) the temporal clustering coefficient had overall moderate test-retest reliability (ICC > 0.40 over a wide range of densities) at both global and subnetwork levels; (2) female subjects showed significantly higher temporal clustering coefficient than males at both global and subnetwork levels, in particular within the default-mode and subcortical regions; (3) temporal clustering coefficient of the subcortical subnetwork was negatively correlated with age in young adults. Our findings suggest that temporal clustering coefficient is a reliable and reproducible approach for the identification of individual differences in brain function, and provide evidence for sex and age effects on human brain dynamic connectome.

## 1. Introduction

Over the past decade, functional magnetic resonance imaging (fMRI)-based brain graphs provide an effective way to modeling the human brain connectome (Bullmore and Bassett, 2011; Cao et al., 2014). A number of topological properties, such as the clustering coefficient and local efficiency that quantify the functional segregation and specialization of brain network (Rubinov and Sporns, 2010; Sporns, 2013), have been widely applied to both healthy and psychiatric populations (Liu et al., 2008; Suo et al., 2018; Tian et al., 2011; Zhang et al., 2011).

Conventional fMRI-based brain graphs are constructed based on the assumption that functional connectivity (FC) patterns are stationary over the period of data acquisition. Recently, however, it has been demonstrated that FC in the brain fluctuates over time, implying that important information may be ignored when describing the brain connectivity in a static manner (Chang and Glover, 2010; Hutchison et al., 2013b). Thus, the investigation of the dynamic functional connectivity (dFC) has rapidly developed which provides new insight into the brain’s dynamical functions (Huang et al., 2021; Hutchison et al., 2013a; Long et al., 2020b; Preti et al., 2017; Valsasina et al., 2019). A commonly used approach for this topic is to model the brain as a multilayer dynamic network (or “temporal network”) in a graph theory-based framework, where a dynamic graph is consisted of multiple time-ordered layers, and each layer corresponds to a “snapshot” of the brain’s functional organization at a particular time point (Boccaletti et al., 2014; De Domenico, 2017; Pedersen et al., 2018). This is an explicit modelling framework that allows us to track where and when entities in a brain network transit from different subsystems, and has greatly enhanced our understanding of the human brain function (Pedersen and Zalesky, 2021).

In the model of dynamic graphs, spatio-temporal properties of the brain network such as “temporal clustering” (Ding et al., 2020; Long et al., 2020a; Ren et al., 2017; Sizemore and Bassett, 2018) and “temporal efficiency” (Fam et al., 2020; Long et al., 2020a; Sizemore and Bassett, 2018; Sun et al., 2019) can be evaluated. These measures are derived from the definitions of analogous metrics for conventional static networks. Specially, the “temporal clustering coefficient”, which is also named “temporal correlation coefficient” by some researchers, quantifies the average topological overlap between any two consecutive layers (i.e. neighboring time points) of the connections in a dynamic brain network (Long et al., 2020a; Nicosia et al., 2013; Sizemore and Bassett, 2018; Tang et al., 2010). Compared with previous approaches quantifying standard deviations/average dissimilarities of dFC patterns across all layers (Long et al., 2020b; Marusak et al., 2017; Zhang et al., 2018; J. Zhang et al., 2016), the temporal clustering coefficient would thus serve as a more direct measure of a network’s tendency to be stable over time. Some recent studies have suggested potential (patho)physiological correlates of this measure: for example, significantly decreased temporal clustering coefficient of brain networks was found to be associated with greater cognitive workloads under complex task conditions in healthy subjects (Ren et al., 2017) and individuals with major depressive disorder during rest (Long et al., 2020a). These reports highlight the potential of temporal clustering coefficient as a novel biomarker in predicting alterations in brain functions in both normal and pathological conditions.

To date, however, at least two empirical questions remain to be addressed before we can use temporal clustering coefficient for large-scale clinical studies. The first question relates to its test–retest reliability; a measure with limited test-retest stability would constrain its clinical utility in longitudinal studies. While reliabilities of some dynamic brain network metrics such as temporal efficiency (Dai et al., 2016) and flexibility (Yang et al., 2021) have been previously estimated and they were thought to be reliable, little has been known thus far in terms of the reliability of temporal clustering coefficient, an index that allows direct inferences on topological stability over time. The second question refers to as whether measures of temporal clustering coefficient would be influenced by sex and age, two critical demographical variables in terms of clinical neuroscience research (Douw et al., 2011; Tian et al., 2011; C. Zhang et al., 2016).

In the present study, we aimed to address the above questions using a publicly available resting-state fMRI (rs-fMRI) dataset from the Human Connectome Project (HCP) (Van Essen et al., 2013) to investigate: (1) test-retest reliability of temporal clustering coefficient; and (2) possible sex- and age-related effects on temporal clustering coefficient. In order to assess the stability of the findings, the derived results were further examined using different analytic pipelines (e.g., the window lengths). For comparative purposes, clustering coefficients for conventional static brain networks were also calculated and compared to the findings in the dynamic networks (e.g., the clustering coefficient (Rubinov and Sporns, 2010)).

## 2. Methods

### 2.1 Datasets and data preprocessing

In line with a previous study (Ji et al., 2019), the analyzed sample consisted of 337 healthy subjects from the “S1200” release of the HCP dataset (Van Essen et al., 2013) with no family relations. All participants were 22 to 37 years old with an average age of (28.61 ± 3.69) years; there were 157 males and 180 females. Each subject underwent four separate rs-fMRI runs during two days, with a repetition time (TR) of 0.72 seconds and a length of 1200 time points for each run. The rs-fMRI data of each run was preprocessed via HCP convention including the ICA-FIX denoising steps, which was thought to be a more selective and effective approach to remove non-neural spatio-temporal components of fMRI signals compared with conventional global signal regression (Glasser et al., 2019, 2018, 2013; Marcus et al., 2013). Further details about the sample information and preprocessing schemes can be found in previous publications (Glasser et al., 2018, 2013; Ji et al., 2019; Marcus et al., 2013).

### 2.2 Dynamic Brain Networks and Temporal Clustering Coefficient

A dynamic network is comprised of a set of nodes and connections between nodes, in which the connections change over time (Holme and Saramäki, 2012; Sizemore and Bassett, 2018). Here, we defined the nodes in brain networks by two different parcellation atlases: 1) the Automated Anatomical Labeling (AAL) atlas (Tzourio-Mazoyer et al., 2002) which subdivides the brain into 90 regions of interest (ROIs) and 2) the Power functional atlas (Power et al., 2011) with 264 ROIs. Both these two atlases have been proved to be valid (Cao et al., 2014) and widely applied in various fMRI studies (Cao et al., 2021, 2018; Long et al., 2019; Tan et al., 2020). All following analyses were repeatedly performed using the two parcellation schemes separately. For each rs-fMRI run of each subject, a multilayer dynamic brain network was then constructed by the sliding-window approach (Laumann et al., 2017; Long et al., 2020a; Reinen et al., 2018) and the temporal clustering coefficient was computed, which was summarized below (also see **Figure 1**):

**Figure 1.**
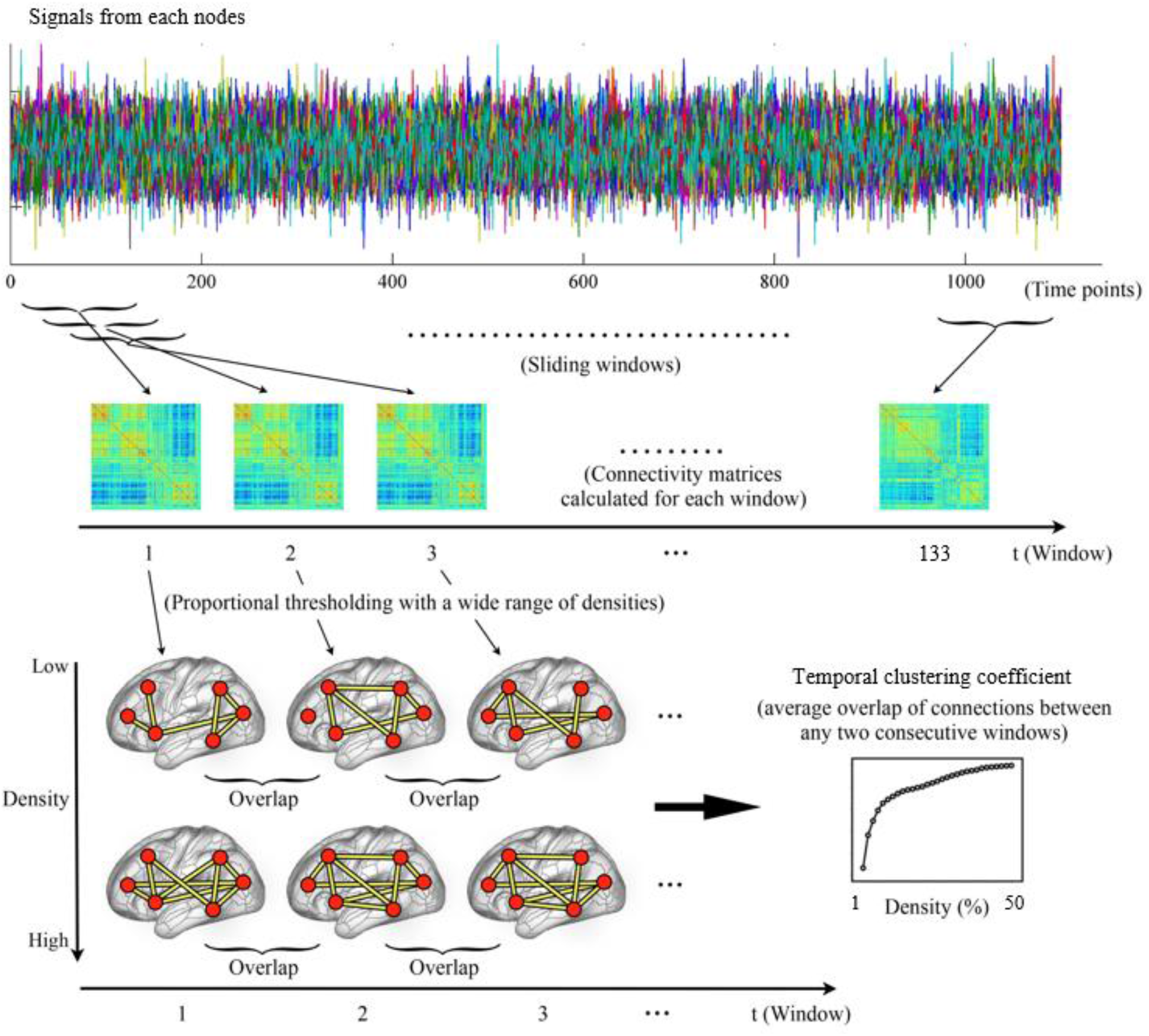
The steps for constructing dynamic brain networks and calculating temporal clustering coefficient (refer to the **Methods** for details).

1. Sliding-windows: firstly, the mean rs-fMRI signals were extracted from each of the 90 or 264 nodes, which were then divided into a series of successive and partly overlapping time windows. Here, a fixed window width of 139 TRs (100.08 seconds) and a window sliding step length of 8 TRs (5.76 seconds) were used in the primary analyses, dividing the whole scan into a total of 133 windows. Note that selections of such parameters were based on recommendations in previous works to balance the quality of connectivity estimation and computational complexity (Long et al., 2020a; Sun et al., 2019; Zalesky and Breakspear, 2015), and effects of changing the window width or sliding step length were also investigated in the latter part of this study. Within each window, the dFC strengths between all pairs of nodes were computed using Pearson correlations, resulting in a 90 × 90 or 264 × 264 connectivity matrix.
2. Proportional thresholding: the temporal clustering coefficient is currently defined in binary networks (Sizemore and Bassett, 2018), which are often obtained by thresholding the networks to preserve only the strongest connections between nodes (Bullmore and Sporns, 2009). In human brain functional connectomics, this was typically achieved by applying a strategy of proportional thresholding (Achard and Bullmore, 2007; Song et al., 2014; van den Heuvel et al., 2017). Here, we applied proportional thresholding on the dFC matrices of each window by preserving only a particular proportion, which is generally called the “density” (Bullmore and Sporns, 2009; Jalili, 2016), of the strongest connections between all possible pairs of nodes. The thresholding was applied with a wide range of densities ranging from 1% to 50% with an increment interval of 1% to ensure that results would not be biased by a single threshold (Bilbao et al., 2018; Cao et al., 2019b). After thresholding, all the preserved connections were assigned a value of 1, and those not surviving the threshold were assigned a value of 0. As a result, a multilayer dynamic network *G* = (*G*_*t*_)_*t* = 1, 2, 3, …, 133_, where *G*_*t*_ is a binary network representing the “snapshot” of dFC pattern within the *t*th time window, was acquired at each density level.
3. Temporal clustering coefficient: at each density, the temporal clustering coefficient was calculated to quantify the overall probability for all connections in the network to persist over consecutive time points. The *nodal temporal clustering coefficient* was firstly computed for each node, as defined by the average overlapping of its connected neighbors between any two consecutive time windows. Briefly, let a_*ij*_ (*t*) =1 if node *i* and node *j* are connected (being neighbors) within the *t*th time window, and a_*ij*_ (*t*) = 0 if they are not. The nodal temporal clustering coefficient of node *i* (*C*_*i*_) was then computed as

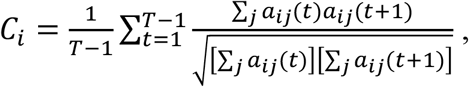

where *T* is the total number of time windows and equals to 133 here (Sizemore and Bassett, 2018; Tang et al., 2010). After that, the global temporal clustering coefficient was obtained by averaging nodal temporal clustering coefficients of all the 90 or 264 nodes (Sizemore and Bassett, 2018). Both the nodal and global temporal clustering coefficients range from 0 to 1, and higher values indicate higher tendencies for the connections to persist across multiple time windows.
4. Subnetwork-level temporal clustering coefficients: according to previous published work (Cao et al., 2019a; Long et al., 2019; Mohr et al., 2016; Power et al., 2011), ROIs in both the two parcellation schemes can be assigned into 9 subsystems including the default-mode, salience, visual, subcortical, auditory, frontoparietal, cinguloopercular, sensorimotor and attention subnetworks. Therefore, we further computed the temporal clustering coefficients for each of these individual subsystems, by averaging the nodal temporal clustering coefficients of all ROIs belonging to that subnetwork.

### 2.3 Static Network Metrics for Comparison

For exploratory comparative purposes, two analogous metrics in conventional static networks, the clustering coefficient and local efficiency (Rubinov and Sporns, 2010) were computed in the binary static brain networks constructed by the whole rs-fMRI scans of each run. Here, they were computed at all densities ranging from 1% to 50%, and at both the global and subnetwork (by averaging the nodal clustering coefficients or nodal local efficiencies of all nodes belonging to particular subnetworks) levels, too.

### 2.4 Test-retest Reliability

As in previous studies (Braun et al., 2012; Cao et al., 2014; Du et al., 2015; Jing et al., 2018; Yang et al., 2021), the test-retest reliability of temporal clustering coefficient was evaluated by the intraclass correlation coefficient (ICC) utilizing a two-way mixed, single measure model (ICC (3, 1)) (Shrout and Fleiss, 1979) as

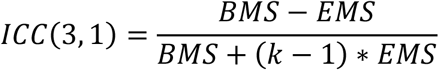

where *BMS* is the between-subjects mean square, *EMS* is the error mean square and *k* is the number of repeated rs-fMRI runs (equals to 4 here). The ICCs were separately calculated at all densities, and at both the global and subnetwork levels. For comparative purposes, ICCs of the two static network metrics (i.e., the clustering coefficient and local efficiency) were also calculated.

The overall ICCs of all metrics were further obtained by averaging across all densities and reported descriptively (Braun et al., 2012; Deuker et al., 2009). According to overall ICCs, the reliabilities of different metrics could be classified into four levels: poor (ICC < 0.4), moderate (0.4 ≤ ICC < 0.6), good (0.6 ≤ ICC < 0.75) and excellent (ICC > 0.75) (Cicchetti, 1994; Roach et al., 2019; Xiang et al., 2021).

### 2.5 Sex- and age-related effects

The possible sex- and age-related effects in temporal clustering coefficients were investigated as follows: 1) for each subject, the temporal clustering coefficient at each density was obtained by averaging all the four rs-fMRI runs; 2) the temporal clustering coefficients were then compared between the male and female subjects using a repeated-measures analysis of covariance (ANCOVA) model in which density (from 1% to 50%) was included as within-subject factor and gender as between-subject factor, covarying for age, education and head motion (as measured by the mean framewise displacement (FD) (Power et al., 2012) across all the four runs); 3) the associations between age and temporal clustering coefficients were investigated by partial Spearman correlations adjusted for sex, education and head motion, where temporal clustering coefficients were averaged over all densities before the correlation analyses. All comparison and correlation analyses were performed at both the global and subnetwork levels. The results were Bonferroni corrected for multiple tests at the subnetwork level and considered significant when corrected *p* < 0.05. For comparative purposes, sex- and age-related effects in the clustering coefficient and local efficiency were also investigated by the same methods.

### 2.6 Data and code availability statement

The “S1200” release of HCP dataset is publicly available at the HCP website (https://www.humanconnectome.org/study/hcp-young-adult). The temporal clustering coefficient was computed based on a publicly-available MATLAB toolbox (Sizemore and Bassett, 2018); the codes can be found at: https://github.com/Yicheng-Long/dynamic_graph_metrics. The clustering coefficient and local efficiency were computed using the Brain Connectivity Toolbox (http://www.brain-connectivity-toolbox.net) (Rubinov and Sporns, 2010).

## 3. RESULTS

### 3.1 Test-retest Reliabilities

As shown in **Figure 2A**, the ICC values of all metrics varied across different densities: the ICC of temporal clustering coefficient varied in the ranges of 0.298-0.565 and 0.340-0.545; the ICC of clustering coefficient varied in the ranges of 0.164 to 0.589 and 0.308 to 0.567; the ICC of local efficiency varied in the ranges of 0.150-0.468 and 0.342-0.497, when using the AAL and Power atlases, respectively.

**Figure 2.**
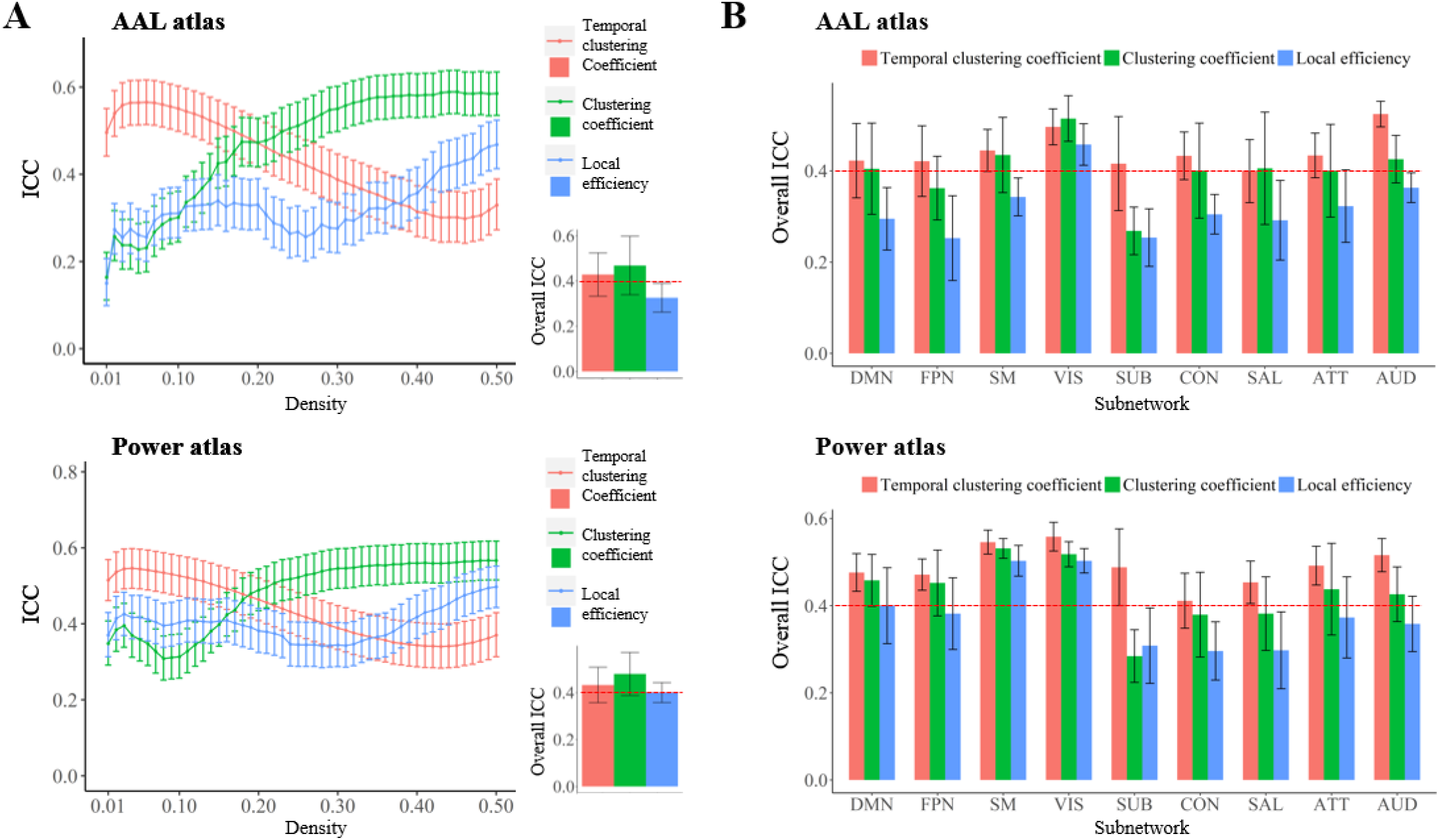
(**A**) Mean values of the intraclass correlation coefficient (ICC) at each density, and the overall ICCs obtained by averaging across all densities (the lower right corner) for each metric at global level. (**B**) The overall ICCs for each metric at subnetwork level. The error bars indicate 95% confidence intervals for ICCs or standard deviations for overall ICCs. ATT, attention subnetwork; AUD, auditory subnetwork; CON, cinguloopercular subnetwork; DMN, default-mode subnetwork; FPN, frontoparietal subnetwork; SAL, salience subnetwork; SM, sensorimotor subnetwork; SUB, subcortical subnetwork; VIS, visual subnetwork.

When averaged across all densities, the overall ICC values of all metrics were mostly in the range of 0.40-0.59, which suggests a moderate test-retest reliability: the temporal clustering coefficient showed an overall ICC of 0.429 ± 0.096 (SD)/ 0.432 ± 0.075 (SD) when using the AAL/Power atlas; the clustering coefficient showed an average ICC of 0.469 ± 0.129 (SD)/ 0.480 ± 0.092 (SD) when using the AAL/Power atlas; the local efficiency showed an average ICC of 0.326 ± 0.063 (SD)/0.400 ± 0.043 (SD) when using the AAL/Power atlas (**Figure 2A**).

Temporal clustering coefficients of each subnetwork were also calculated and **Figure 3** shows their correlations with the global temporal clustering coefficient. As shown in **Figure 2**, overall ICCs of the temporal clustering coefficients were found to be higher than 0.4 for most subnetworks, which suggests a moderate test-retest reliability. Meanwhile, overall ICCs of the clustering coefficient and local efficiency for most subnetworks were found to be relatively lower than those of the temporal clustering coefficients for the same subnetworks (**Figure 2**).

**Figure 3.**
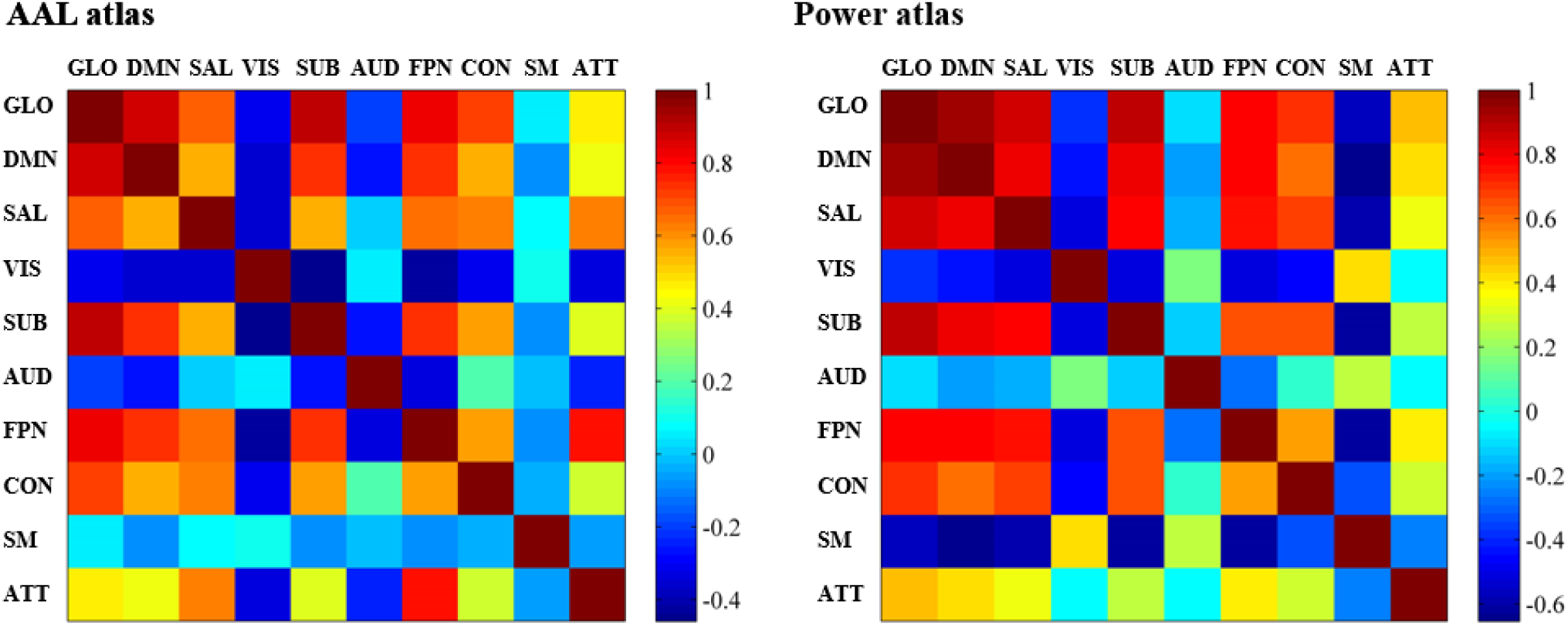
Pearson correlation coefficients between the global temporal clustering coefficient and temporal clustering coefficients of each subnetwork. ATT, attention subnetwork; AUD, auditory subnetwork; CON, cinguloopercular subnetwork; DMN, default-mode subnetwork; FPN, frontoparietal subnetwork; GLO, global; SAL, salience subnetwork; SM, sensorimotor subnetwork; SUB, subcortical subnetwork; VIS, visual subnetwork.

### 3.2 Sex- and age-related effects

As shown in **Figures 4-5**, female subjects showed a significantly higher global temporal clustering coefficient than male subjects (*F* = 13.220/18.280, *p* = 3.21×10^−4^/2.50×10^−5^ when using the AAL/Power atlases). At the subnetwork level, significantly high temporal clustering coefficients in female subjects were found within the default-mode and subcortical subnetworks (Bonferroni-corrected *p* < 0.05), while no significant gender differences were found in other subnetworks (corrected *p* > 0.05). All the above results were robustly found with both the two parcellation schemes (AAL/Power atlases).

**Figure 4.**
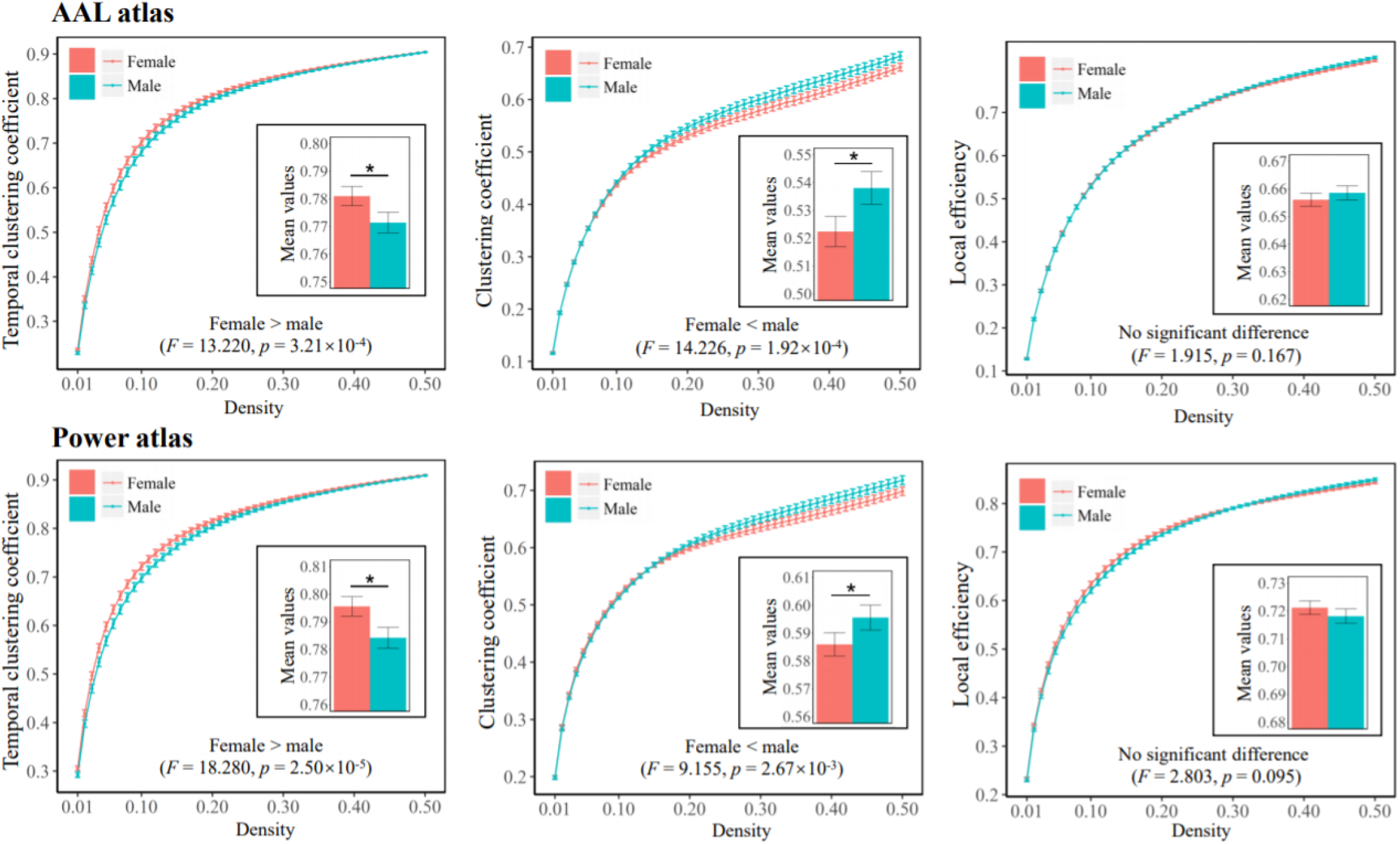
Comparisons of each metric at global level between the male and female subjects. The mean values for each metric obtained by averaging across all densities (0.01 to 0.50) are also presented. The “*” indicates a significant difference and error bars indicate 95% confidence intervals.

**Figure 5.**
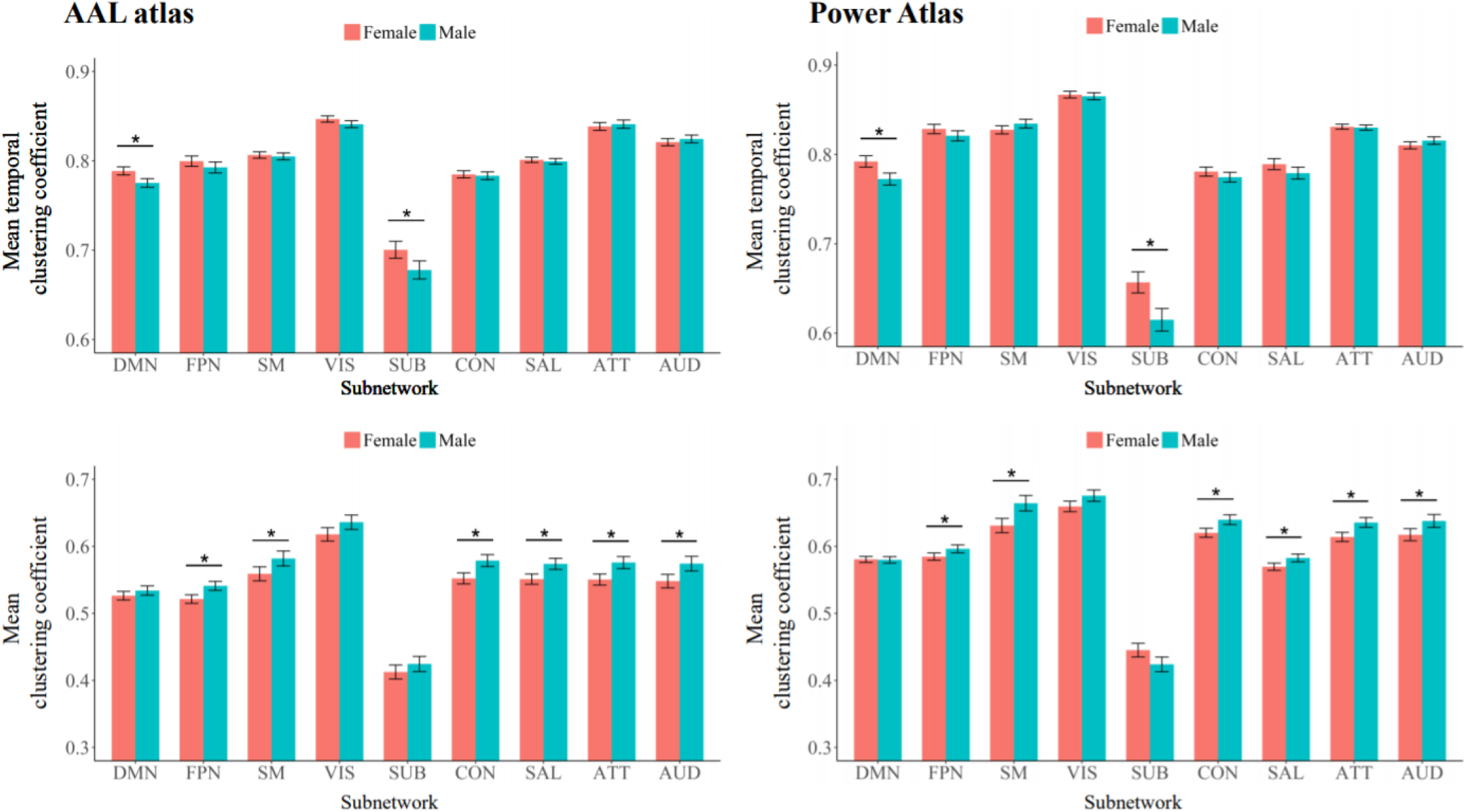
Comparisons of subnetwork-level temporal clustering coefficients and clustering coefficients between the male and female subjects. The mean values for each metric obtained by averaging across all densities (0.01 to 0.50) are presented here. The “*” indicates a significant difference (with Bonferroni-corrected *p* < 0.05) and error bars indicate 95% confidence intervals. ATT, attention subnetwork; AUD, auditory subnetwork; CON, cinguloopercular subnetwork; DMN, default-mode subnetwork; FPN, frontoparietal subnetwork; SAL, salience subnetwork; SM, sensorimotor subnetwork; SUB, subcortical subnetwork; VIS, visual subnetwork.

Results of the age effects are presented in **Figures 6-7**. At the global level, the results showed a positive relationship between age and temporal clustering coefficient which was, however, not significant based on the Power parcellation (Spearman’s rho = 0.131/0.087, *p* = 0.016/0.113 when using the AAL/Power atlases). At the subnetwork level, a significant positive correlation was found between age and temporal clustering coefficient of the subcortical subnetwork (Spearman’s rho = 0.174/0.173, uncorrected *p* = 0.001/0.001, Bonferroni corrected *p* = 0.012/0.013 when using the AAL/Power atlases) with both the two parcellation schemes. As for static network metrics, at the global level, significant negative correlations were found between age and clustering coefficient (Spearman’s rho = -0.166/-0.128, *p* = 0.002/0.019 when using the AAL/Power atlases), as well as between age and local efficiency (Spearman’s rho = -0.194/-0.109, *p* = 0.0003/0.046 when using the AAL/Power atlases); however, no significant correlations were found at the subnetwork level (corrected *p* > 0.05 for all subnetworks).

**Figure 6.**
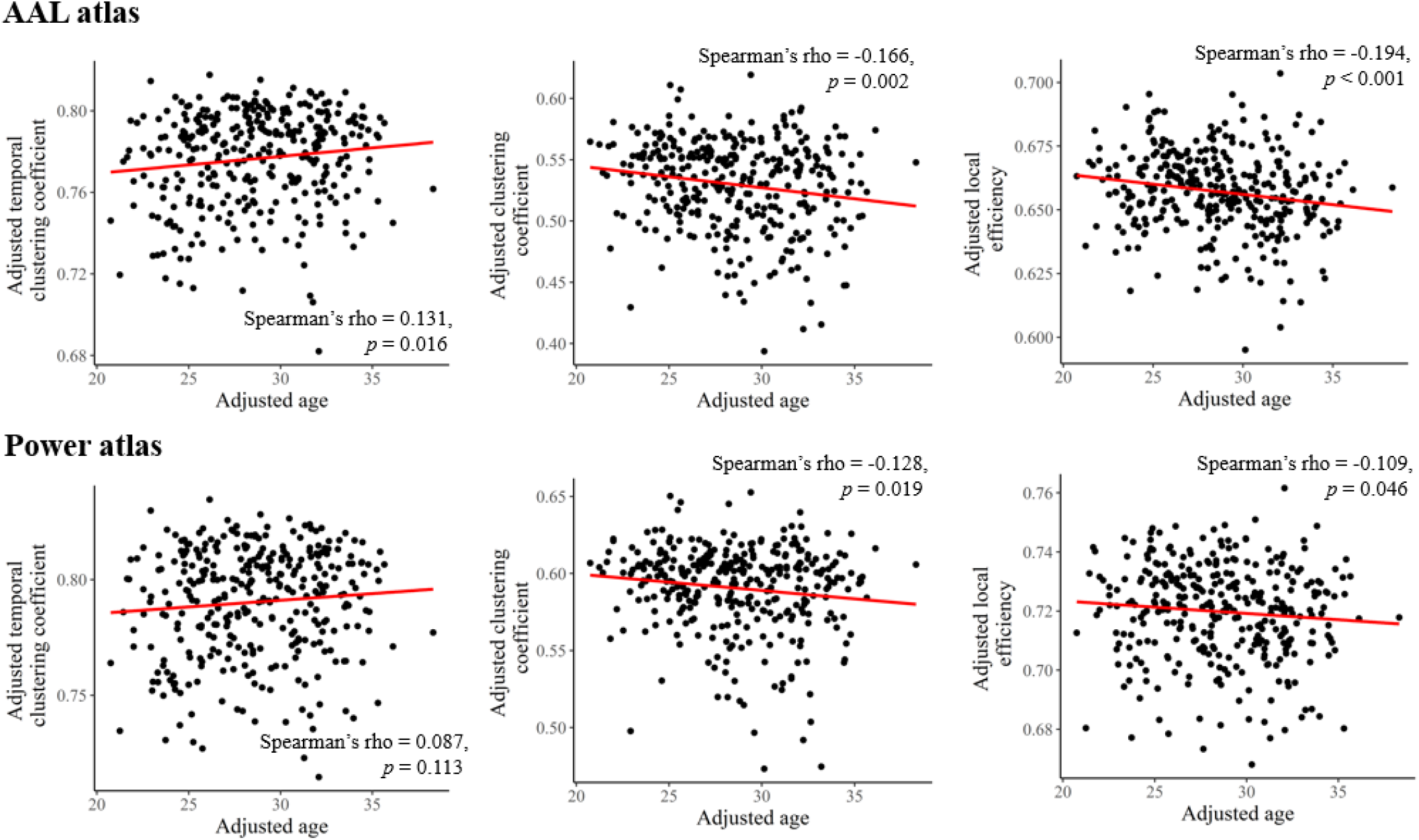
Results of partial Spearman correlations between age and each brain network metric at global level, after adjusting for sex, education and head motion. All metrics were averaged across all densities (0.01 to 0.50) before the correlation analyses.

**Figure 7.**
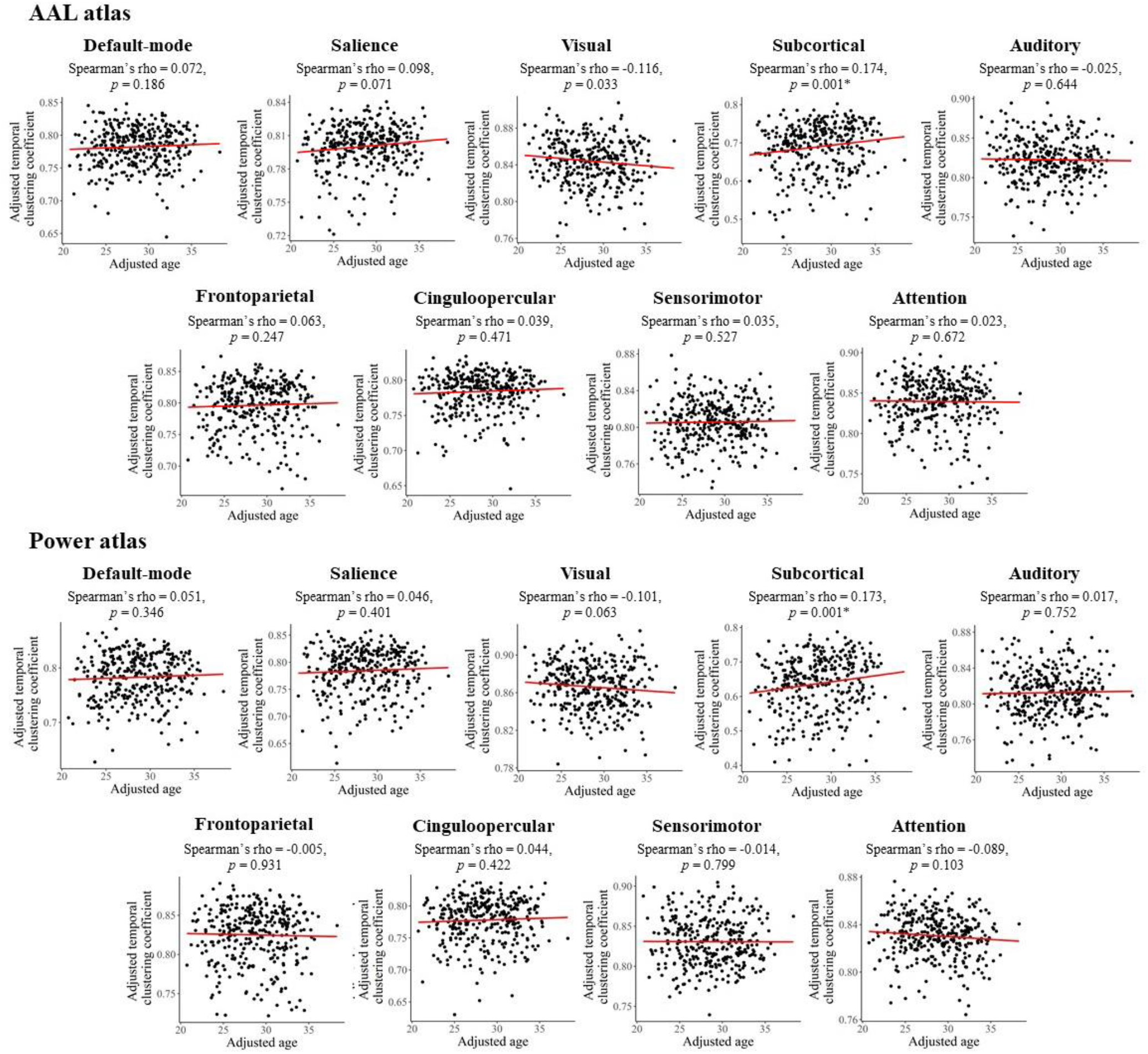
Results of partial Spearman correlations between age and temporal clustering coefficients of each subnetwork, after adjusting for sex, education and head motion. The temporal clustering coefficients were averaged across all densities (0.01 to 0.50) before the correlation analyses. Uncorrected *p* values are presented and the “*” indicates a significant correlation with Bonferroni-corrected *p* < 0.05.

## 4. Discussion

In this study, we investigated test-retest reliability and sex- and age-related effects on a newly introduced dynamic graph-based metric called temporal clustering coefficient in human functional brain networks. Overall, our results support three main conclusions. First, the temporal clustering coefficient showed moderate test-retest reliability at both global and subnetwork levels, which is comparable to those of analogous conventional static brain network metrics (e.g., clustering coefficient and local efficiency). Second, significantly higher temporal clustering coefficients in female subjects than males were found at global level and within the default-mode and subcortical regions. Third, a significant positive relationship was found between age and temporal clustering coefficient at the subnetwork level within the subcortical regions.

The first major contribution of this study is the evaluation of test-retest reliability of temporal clustering coefficient in human brain network for the first time to our knowledge. The temporal clustering coefficient (also named temporal correlation coefficient) has previously been used to characterize the persistence of connections over time in time-varying systems such as the public transportation system (Galati et al., 2013), trade network (Büttner et al., 2016), and the human brain functional networks (Long et al., 2020a; Ren et al., 2017; Sizemore and Bassett, 2018). However, it is unknown about the reliability of this approach when applying to human brains. In this study, our results revealed fair test-retest reliability (overall ICC > 0.4) for temporal clustering coefficients in both the whole brain and individual large-scale subnetworks (**Figure 2**). Our results thus support that the temporal clustering coefficient would be a reliable metric to characterize the brain functional dynamics.

The second important finding of our study is a reproducible sex difference in temporal clustering coefficient. Using two different parcellation schemes, a significantly higher temporal clustering coefficient was consistently observed in female subjects than males, for both the entire brain (**Figure 4**) and the default-mode and subcortical systems at subnetwork level (**Figures 5**). A higher “temporal clustering” indicates a higher average overlap of the network structures between consecutive time points (Sizemore and Bassett, 2018). Our results thus, intriguingly, suggest a higher trend for dFC patterns to be persistent over time in the brains of females than those of males. To our knowledge, investigations on differences in dFC between males and females are still limited, with various methods employed and inconsistent results (de Lacy et al., 2019; Mao et al., 2017; Menon and Krishnamurthy, 2019; Yaesoubi et al., 2015). For instance, by comparing the “flexibility” as defined by switching frequency of the network modular community structures, an earlier study has reported higher flexibility in females than males within regions of the default-mode network, thus suggesting a lower temporal stability of dFC patterns in females (Mao et al., 2017). Another research explored the brain dynamics by using a state-clustering algorithm andsuggested that females switch connectivity states less frequently and spend more time in the state where dFC within the default-mode network is strong (de Lacy et al., 2019). Here, our results were in line with the latter one. Considering that our results were obtained from a relatively large sample and confirmed with different parcellation schemes, the present study may provide futher evidence that female subjects have temporally more stable functional brain organizations than males, especially within the default-mode and subcortical regions. Notably, alterations in temporal stability of dFC within these systems have been assiociated with a number of common mental diseases such as major depressive disorder (Long et al., 2020a; Wise et al., 2017) and bipolar disorder (Nguyen et al., 2017), which have marked gender biases in incidences (Seedat et al., 2009). Therefore, it is hypothesized altered dFC may play important roles in such gender differences, which can be further investigated in future studies.

Besides the sex differences, our results also showed a significant positive correlation between age and temporal clustering coefficient of the subcortical subnetwork (**Figure 7**). There were some previous studies based on different methodologies reporting that temporal variability of dFC “states” involving the subcortical regions were negatively correlated with age (Xia et al., 2019). Therefore, out results were in line with the previous report, and may suggest an increasing temporal stability (decreasing temporal variability) of the subcortical dFC patterns during the process of aging. It was noteworthy that apart from the subcortical regions, significant age effects on dFC variability within some other brain systems such as the default-mode cortices have also been reported (Park et al., 2017; Qin et al., 2015) but they were not observed in this study. One possible reason is that age range of the current sample is relatively narrow (22 to 37 years old). In future studies, it might be necessary to confirm if similar age effects on the temporal clustering coefficient would exist over a broader range of age, or even lifespan.

In recent years, there have been some efforts to understand how the brain’s functional organization fluctuates over time, pointing out that it is temporally changed in specific manners (Ma and Zhang, 2018; Vidaurre et al., 2017). It has been suggested that, for instance, the dFC patterns of brain do not fluctuate randomly but tend to persist over some periods of time (Allen et al., 2014; Tang et al., 2010). Based on this important property, the activity of human brains during rest can be temporally clustered into discrete short periods or “connectivity states”, during which the dFC patterns remain quasi-stable over tens of seconds (Damaraju et al., 2014; Leonardi et al., 2013). Furthermore, alterations related to such connectivity states have been reported in various psychiatric disorders (Damaraju et al., 2014; Rashid et al., 2014; Reinen et al., 2018; Zhi et al., 2018). Despite these accumulating findings, little has been known as whether there is a reliable measure to quantify the tendency for dFC patterns to persist over time, and to detect the inter-individual differences in such tendencies. Here, we propose that the measure of temporal clustering coefficient may be a suitable indicator of temporal persistence of the brain’s functional structures, and could potentially be used to predict alterations in brain functions given its acceptable reliability and high reproducibility as shown by our results.

For exploratory comparative purposes, we repeated the whole analyses on two analogous static network metrics including the clustering coefficient and local efficiency. It was found that temporal clustering coefficient had a close overall ICC compared with these two metrcis at global level (**Figure 2A**). Moreover, the temporal clustering coefficient had relatively higher ICCs than those of clustering coefficient and local efficiency for most subnetworks at local level; and specially, the difference is most pronounced for the subcortical subnetwork (**Figure 2B**). As a possible explanation, we consider this is due to the fact that subcortical structures such as the thalamus and basal ganglia relay and modulate information passing to different areas of the brain; their dFC patterns are most variable over time since they may connect with different brain regions and be involved in different functional communities/modules at different times (Long et al., 2020b; J. Zhang et al., 2016). Therefore, it might be more precise to characterize these regions by dFC rather than conventional static FC. As for sex- and age-related effects, several significanlt results were obtained for both the clustering coefficient and local efficiency, which are in line with some previous research (**Figures 4-6**) (Douw et al., 2011; Tian et al., 2011). Importantly, it was shown that compared with clustering coefficient and local efficiency, the temporal clustering coefficient showed significaint sex/age-related effects within different subnetworks, indicating that they may reflect different aspects of brain function. This may partly support the opinion that dFC can capture important information ignored by static methodology (Chang and Glover, 2010; Hutchison et al., 2013b), and further highlight the value of temporal clustering coefficient.

There are some limitations of this study and possible future directions to be noted. First, the temporal clustering coefficient is currently defined based on binary graphs (Sizemore and Bassett, 2018), which ignore much important information contained in the connection weights (Cao et al., 2014; Rubinov and Sporns, 2010). The definition of temporal clustering coefficient may be expanded to weighted graphs in future studies to deal with such weakness. Second, while we performed all analyses using two validated atlases for node definition and a commonly used rs-fMRI experiment, it may be necessary to confirm our results with some other parcellation schemes and tasks. Last, exploring possible alterations in temporal clustering coefficient in psychiatric populations, rather than only in healthy participants, may provide much more important clinical implications for the brain functional dynamics.

## 5. Conclusions

In summary, in this study, we investigated the test-retest reliability of a novel dynamic graph-based metric called temporal clustering coefficient, and possible sex- and age-related effects in such a measure. The results demonstrated fair test-retest reliability of temporal clustering coefficient at both global and subnetwork levels, suggesting that it would be a reliable approach to find meaningful individual differences in brain function. Moreover, reproducible higher global temporal clustering coefficient was found in females than males, in particular the default-mode and subcortical subnetworks at local level, suggesting a temporally more stable functional brain organization in females. At local level, significant positive correlation was also found between age and temporal clustering coefficient of the subcortical subnetwork, which may suggest increased temporal stability of subcortical dFC patterns along age. These results may serve as an initial guide for evaluating human brain functional dynamics.

## ACKNOWLEDGEMENTS

This work was supported by the Natural Science Foundation of Hunan Province, China (grant numbers 2021JJ40851 and 2021JJ40835), the Changsha Municipal Natural Science Foundation (kq2014238) and National Natural Science Foundation of China (grant number 82071506). Data were provided by the Human Connectome Project, WU-Minn Consortium (Principal Investigators: David Van Essen and Kamil Ugurbil; 1U54MH091657) funded by the 16 NIH Institutes and Centers that support the NIH Blueprint for Neuroscience Research; and by the McDonnell Center for Systems Neuroscience at Washington University. The author Yicheng Long would like to thank Dr. Alan Anticevic and Jie Lisa Ji from Yale University for their suggestions in the preparation of this study.

